# Lung at a Glance: an integrative web toolset of lung ontology, imaging and single cell omics

**DOI:** 10.1101/2020.06.19.161851

**Authors:** Yina Du, Weichen Ouyang, Joseph A Kitzmiller, Minzhe Guo, Shuyang Zhao, Jeffrey A Whitsett, Yan Xu

## Abstract

Recent advances in single-cell omics and high-resolution imaging have provided unanticipated data resources for the elucidation of genes underlying the complex biological processes critical for organ formation and function. However, processing and integrating large amounts of single-cell omics and imaging data presents a major challenge for most researchers. There is a critical need for ready-to-use computational tools for data/knowledge integration and visualization. Here we present “Lung-at-a-glance”, an easy-to-use web toolset for visualizing and interoperating complex omics and imaging data, providing an interactive web interface to bridge lung anatomic ontology classifications to lung histology and immunofluorescence confocal images, and cell-type-specific gene expression. “Lung-at-a-glance” contains three interactive components: 1) “Region at a glance”, 2) “Cell at a glance” and 3) “Gene at a glance”. “Lung-at-a-glance” and other newly developed web tools for lung-related data query, integration and visualization are publicly available on LGEA web portal v3 https://research.cchmc.org/pbge/lunggens/mainportal.html.

## INTRODUCTION

LGEA web portal offers user-friendly and comprehensive web-based query functions on extensive bulk, sorted, single-cell transcriptomic and image data from human and mouse lung at different stages of development (1, 2). LGEA web portal has been used by researchers from more than 130 institutes from 52 different countries and has been cited by more than 120 scientific publications. With the continuing support from the LungMAP consortium, we extended the scope of the LGEA web portal database to include proteomics and epigenetic data from lung developmental and lung disease studies. To facilitate utilization and integration of these data resources, we developed “Lung-at-a-glance” to serve as an interactive bridge to connect lung anatomic ontology, histology and lung gene/protein expression data with easy-to-use web interfaces. A number of other newly developed bioinformatics tools for users to query, compare and integrate diverse lung omics datasets are also made available on LGEA new release.

## METHODS

Modern JavaScript-based charting libraries (https://www.highcharts.com), CSS framework of Bootstrap (https://getbootstrap.com) and W3.CSS (https://www.w3schools.com/w3css) are integrated for web design and data visualization. The new features are implemented into the existing LGEA web architecture to improve its stability and modularity, including separation of front-end and back-end, and continuous integration and delivery. Oracle database management system 12c (https://www.oracle.com/database) is used for central data storage and maintenance. UCSC Genome Browser (https://genome.ucsc.edu) is embedded into web application to provide an interactive visualization of epigenetic data.

## RESULTS

### “Lung-at-a-glance”, a featured toolset in LGEA

“Lung-at-a-glance” consists of three interactive components: “Region-at-a-glance”, “Cell-at-a-glance”, and “Gene-at-a-glance”; all designed to provide data access with a single-click on the icons located on the top of “Lung-at-a-glance” homepage (https://research.cchmc.org/pbge/lunggens/tools/lung_at_a_glance.html). The “at-a-glance” toolset is designed to serve as an interactive bridge to connect lung anatomic ontology, histology and immunofluorescence confocal images in Lung Image gallery (https://research.cchmc.org/lungimage/) and lung gene expression data in the LGEA web portal. The toolset provides a collection of comprehensive interrelated data and knowledge resources with a user friendly intuitive and interactive web-interface for data analysis, integration and visualization (**Figure 1**). The anatomic ontology for human and mouse lung was developed by NHLBI LungMAP Consortium, Ontology Subcommittee using Web Ontology Language (OWL 2). The ontology contains ∼ 300 terms for fetal and postnatal lung structures, tissues, and cells which were identified for each species (3). We converted the abstract version of anatomic ontology terms into searchable, clickable and expandable web tree structures, we further incorporated the ontology tree as an interactive component into LGEA. Investigators can navigate the hierarchical structure of the anatomical tree or use the search box to directly locate regions or cells of interest.

**Figure 1.**
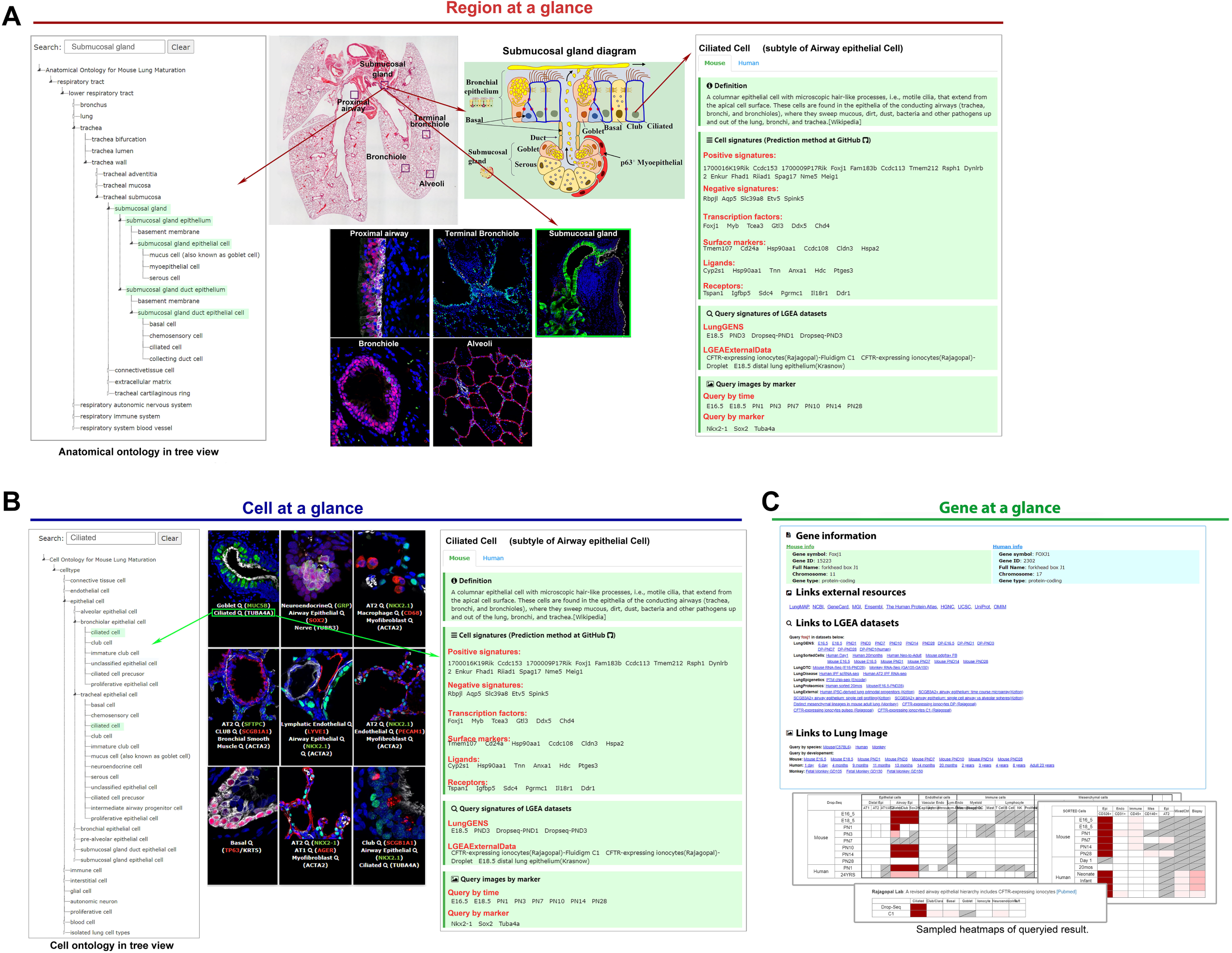
“Lung-at-a-glance” consists of three interactive and interconnected components: “Region at a glance”, “Cell at a glance”, and “Gene at a glance”. (A) “Region at a glance” allows users to search a specific lung region using the interactive ontology tree navigation or clicking the boxes in the lung image to explore cells reside in the selected region. (B) “Cell at a glance” offers a collection of information of queried cell type, including cell definition, cell type specific positive and negative markers, transcription factors, ligand-receptors, and hyperlinks which are mapped to the chosen cell type in all datasets in LGEA web portal and immunofluorescence confocal images of the chosen cell type. (C) “Gene at a glance” allows users to query a gene of interest to obtain associated gene information including hyperlinks to external knowledge base, to expression patterns in LGEA datasets, and to immunofluorescence confocal images. A two-dimensional heatmap by developmental times and cell types was used to summarize expression patterns and statistics of queried gene in all datasets available in LGEA databases.

“Region-at-a-glance” allows users to search a specific lung region using the interactive tree navigation or click one of the annotated lung regions, e.g., “Proximal airway”, “Submucosal gland”, “Bronchiole”, “Terminal Bronchiole”, and “Alveoli” on the H&E staining lung image. Users can explore cells within selected regions using interactive mouse-hover features which are embedded in the anatomical ontology tree, images and diagrams (**Figure 1A**). “Cell-at-a-glance” can be activated by clicking the cell name or image on the “Cell-at-a-glance” page or by clicking a cell of interest in selected regional diagrams on the “Region-at-a-glance” page. With the same interactive web page design pattern, “Cell-at-a-glance” offers a collection of information related to the queried cell type including cell definition, cell type specific positive and negative markers, transcription factors, ligand-receptors predicted by our group (https://github.com/xu-lab/LGEA_Cell_Signature), and hyperlinks to all LGEA web portal datasets and immunofluorescence confocal images related to the chosen cell type. Approximately 40 cell types are available for study in the current “Cell-at-a-glance” (**Figure 1B**). “Gene-at-a-glance” allows users to query a gene of interest and obtain its expression patterns (including protein expression levels if available) and statistics in all LGEA datasets in a 2-D heatmap linking developmental times and cell types (**Figure 1C**). Hyperlinks to external knowledge-bases and immunofluorescence confocal images of relevant cell markers are provided in the “Gene-at-a-glance” page (**Figure 1C**). The three components of “Lung-at-a-glance” are interconnected offering users a one-stop bioinformatics-tool for lung research. For example, investigators can start their search at specific anatomic regions, explore a particular cell type within the region, and identify cell specific markers, ligands-receptors and transcription factors expressed in the cell type of interest across lung developmental stages.

### Diverse datasets and web functionality

In addition to “Lung-at-a-glance”, the LGEA new release represents a significant update of the previous version, expanding to ten functional query panels from three panels in the previous version (1, 2). The analytic tools and functionalities of current LGEA web portal are briefly introduced below (**Figure 2**).

**Figure 2.**
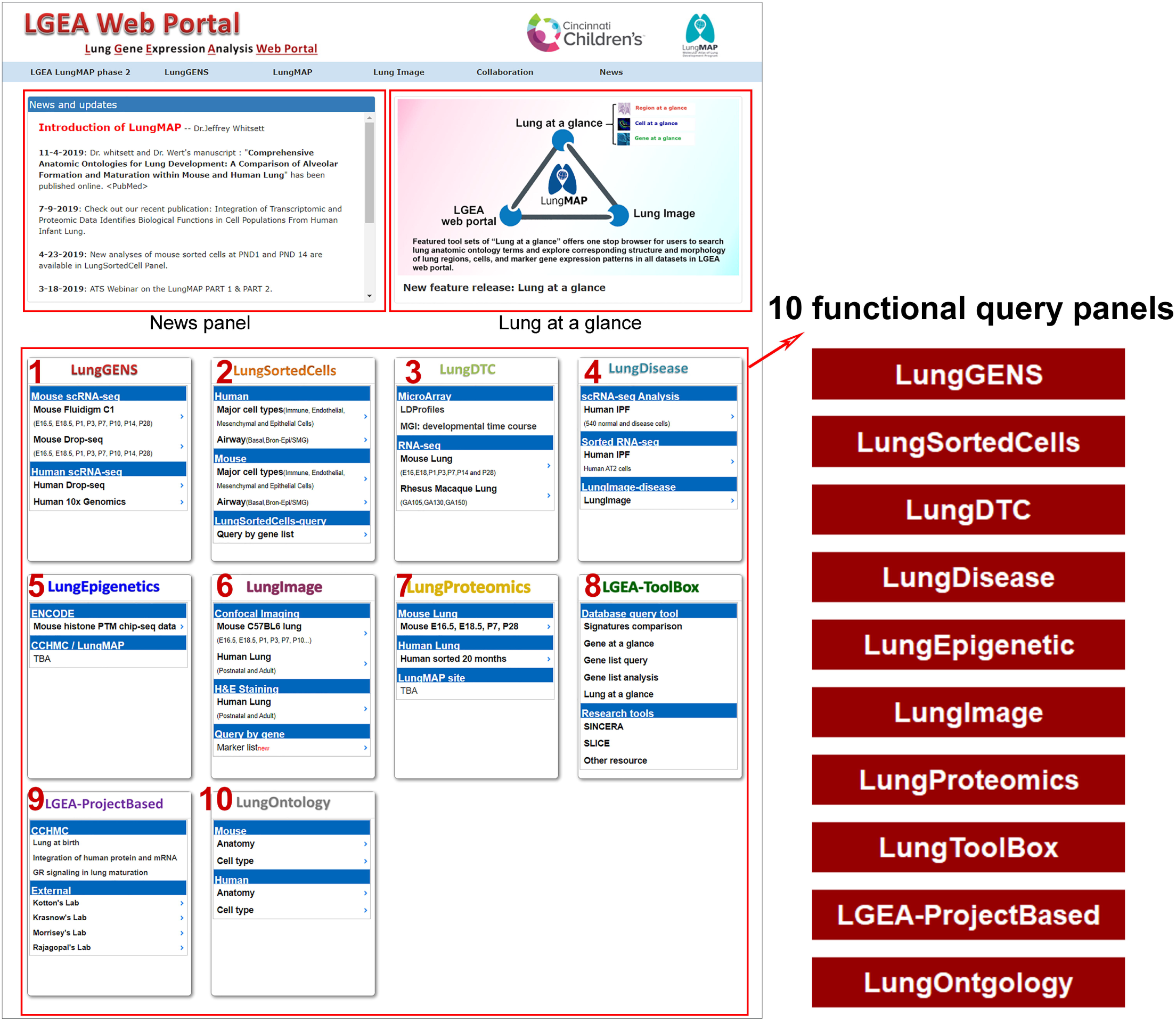
LGEA web portal home page. The new release of LGEA web portal provides query and analytic tools including “LungGENS”, “LungSortedCells”, “LungDTC”, “LungDiseases”, “LungEpigenetics”, “LungImage”, “LungProteomics”, “LungOntology”, “LGEA-ProjectBased” and “LGEA-ToolBox”. Newly developed feature toolset “Lung-at-a-glance” is highlighted on top of the home page.

- The “LungGENS”, “LungSortedCells”, “LungDTC” supports ‘Query by single gene’, ‘Query by gene list’ and ‘Query by cell type” function on single-cell, sorted-cell and whole lung developmental time course data collected from mouse lung tissues from embryonic day 16.5 to postnatal day 28 (eight time points) and human lung samples from neonate, infant, child and adult at various ages.
- The “LungDisease” panel allows investigators to search altered gene expression patterns, disease related signatures and lung disease related images including idiopathic pulmonary fibrosis(4), lymphangioleiomyomatosis (5), cystic fibrosis, bronchopulmonary dysplasia (BPD), primary alveolar microlithiasis, asthma, ABCA3 mutations, TBX4 deletion and lung growth disorder.
- The “LungEpigenetics” panel contains ChIP-seq data during mouse lung development downloaded from ENCODE (https://www.encodeproject.org/). Developmental stages and specific histone modifications associated with the gene of interest are viewed on the embedded UCSC Genome Browser with selected tracks enabled.
- “LungProteomics” provides protein centered query functions based on proteomic data generated at Pacific Northwest National Laboratory (PNNL) in the LungMAP project. Correlations between protein and mRNA expression profiles, and an overview of expression levels of the queried protein/gene expression across all data sets in the LGEA database can be visualized.
- “LGEA-ProjectBased” is designed to introduce and share high impact lung related projects from internal and external research groups.
- The LGEA-ToolBox includes a number of new bioinformatics tools, including “signature comparison”, “gene list query”, and “gene list analysis” to facilitate comparison and integration of datasets to meet the ever-increasing needs of the research community. Users can compare different datasets in LGEA database or enter their gene list of interest to compare cell type specific signature genes provided by LGEA.

## DISCUSSION

Recent advances in single-cell omics have provided increasing insights into the pathogenesis of human disease, including those affecting the lung (6-10). The density of omics data relevant to lung biology is increasing exponentially, providing information that, with suitable integrative analysis, may provide important insights into the processes underlying lung function. The LGEA web portal is designed for intuitive and practical interrogation use of comprehensive “omics” data obtained during normal lung morphogenesis and from lung diseases by research investigators with varying computational training. LGEA new release provides improved interactive, graphical web interfaces for search, visualization, and secondary analyses, in which output can be readily interpreted and downloaded. Featured toolset of “Lung-at-a-glance” offers end-to-end web functions to access and search lung anatomic ontology terms, and to explore the corresponding structure and morphology of tissue regions, cells, and marker gene expression patterns in all datasets in the LGEA database. To our knowledge, this is the first web application connecting anatomic ontology terms to structure and histology images and to single cell expression data. These tools and enriched data resources can be used to enhance hypothesis generation and scientific discovery. The LGEA web portal database will be continually updated with more omics data generated in LungMAP 2 and new query functions will be developed to further enhance data and knowledge interrogation. Data are made available and synchronized at the LungMAP website (https://www.lungmap.net/).

## ACKNOWLEDGEMENTS

The authors acknowledge the support of Dr. Sara Lin (Program Director) and all members of LungMAP research Consortium.

## DATA AVAILABILITY

LGEA web portal v3 are freely available for non-commercial use at http://research.cchmc.org/pbge/lunggens/mainportal.html and data is readily integrated with omics data and lung image data from other research centers at “BREATH” database and display on the LungMAP website (https://www.lungmap.net/).

